# Targeting triple-negative breast cancer with β1-integrin binding aptamer

**DOI:** 10.1101/2022.07.28.501822

**Authors:** Karlis Pleiko, Maarja Haugas, Vadims Parfejevs, Teodors Pantelejevs, Emilio Parisini, Tambet Teesalu, Una Riekstina

## Abstract

Targeted therapies have increased the treatment options for triple-negative breast cancer patients. However, the paucity of targetable biomarkers and tumour heterogeneity have limited the ability of precision-guided interventions to live up to their full potential. As affinity targeting ligands, aptamers show high selectivity towards target molecules. Compared to antibodies, aptamers have lower molecular weight, increased stability during transportation, reduced immunogenicity, and increased tissue uptake. Recently, we reported the discovery of GreenB1 aptamer that is internalized in cultured triple-negative MDA-MB-231 human breast cancer cells. We show that the GreenB1 aptamer specifically targets β1-integrin, a protein previously linked to breast cancer cell invasiveness and migration. Aptamer binds to β1-integrin with low nanomolar affinity. GreenB1 homes in the orthotopic 4T1 triple-negative breast cancer lesions modelled in mice. Our findings suggest potential applications for the GreenB1-guided precision agents for the diagnosis and therapy of triple-negative breast cancer.

## INTRODUCTION

Triple-negative breast cancer (TNBC) accounts for ∼20% of all invasive breast cancer cases. TNBC tumours are negative for expression of human epidermal growth factor receptor 2 (HER-2), progesterone receptor (PR), and estrogen receptor 2 (ER), rendering TNBC resistant to endocrine therapy (1). Combination of chemotherapy followed by surgery is used as a treatment strategy for early TNBC, while chemotherapy is used to treat advanced and metastatic TNBC (2). Treatments with immune-checkpoint inhibitors (ICIs) targeting programmed cell death protein 1 (PD-1) and programmed death-ligand 1 (PD-L1) have augmented the therapeutic choices for patients with PD-L1^+^ TNBC in recent years (3).

Appreciation of the heterogeneity of the TNBC microenvironment (TME), including differences in immunological composition, vascularization, metabolic status, and stromal composition, has resulted in identifying TNBC subtypes with different treatment responses (4). TNBC has at least three subtypes (basal, luminal androgen receptor and mesenchymal). Single-cell sequencing has revealed tumour microenvironment heterogeneity, showing populations of cells typical of cancers with poor outcomes (5) and subtypes based on gene-regulatory networks (6). The development of therapies that target each subtype may increase the number of available treatment options in the future (7). Based on TME differences, tumours can be divided into “hot” tumours, a T cell inflamed cancer phenotype, and “cold” tumours, non T cell inflamed phenotype. Current ICIs are limited to acting on hot tumours (8). Anticancer vaccines, targeted therapies that increase the re-expression of tumour-associated antigens, engineered T cells expressing chimeric antigen receptors (CARs), and other (5) approaches are being studied to promote T cell infiltration, thereby transforming cold tumours into ICI responsive hot tumours. Furthermore, targeted therapy is used against TME cellular components; for example, OximUNO (nanoconjugate of CD206 targeting peptide mUNO with doxorubicin) has shown promise in pre-clinical studies to inhibit breast cancer progression by depleting anti-inflammatory tumour-supporting macrophages (9).

Antibody-drug conjugates (ADCs) have been used successfully as guided precision agents (10). One such ADC, sacituzumab govitecan, composed of antibody targeting trophoblast cell-surface antigen 2 (TROP2) linked to SN-38 (topoisomerase I inhibitor) through a hydrolyzable linker has received FDA approval for the treatment of metastatic TNBC (11). Several other ADCs are undergoing clinical trials and have been reviewed elsewhere recently (12). The bispecific antibody PF-06671008, which targets CD3 on T cells and P-cadherin (CDH3) on tumour cells, is another promising strategy for T-cell recruitment to tumour sites (13). It has been investigated in a phase I clinical trial (ClinicalTrials.gov, NCT02659631) for treatment of advanced solid tumours. However, present treatments do not yet provide optimal therapy options.

Aptamers are short (20-100 nt), single-stranded DNA or RNA oligonucleotides that bind to their target molecules due to a specific three-dimensional structure. Their affinity and specificity are comparable to antibodies; however, aptamers are smaller (6-30 kDa versus 150-180 kDa for antibodies) and can be chemically synthesised, resulting in minimal to no batch-to-batch variability and straightforward scale-up. Aptamers are stable, can be denatured/refolded, show a rapid tissue uptake (14) and low immunogenicity (15). Recently, Kelly et al. have highlighted considerable difficulties in translating aptamers selected under cell-free settings to *in vitro* and *in vivo* studies. Out of 15 reported aptamers against cell surface targets, 5 demonstrated receptor-specific activity on cells *in vitro*, and only one (anti-hTfR aptamer Waz) was able to target tumours *in vivo* (16).

Target-specific aptamers have been utilised to create tools for detecting circulating targets (circulating tumour cells, proteins, extracellular vesicles) (17, 18), aptamer-targeted vesicles or nanoparticles that improve medication delivery (19, 20) and fluorescent RNA-based biosensors for metabolite detection (21).

Selections on TNBC-related proteins or on cultured TNBC cells have identified multiple aptamers. Epidermal growth factor receptor (EGFR) (22–25), platelet-derived growth factor receptor β (PDGFRβ) (26–28), nucleolin (29–35), CD133 (36), CD44 (37, 38), epithelial cell adhesion molecule (EpCAM) (39, 40), CD49c (41) and Tenascin-C (42) binding aptamers have shown the potential of selective delivery of therapeutic agents to TNBC *in vitro* and *in vivo*.

Here we show that TNBC cell line selective aptamer GreenB1 binds to β1-integrins and is internalized in cells. We also demonstrate that upon systemic administration, GreenB1 homes to the orthotopic TNBC lesions modelled in mice.

## MATERIALS AND METHODS

### *In vitro* cell cultures

Estrogen and progesterone receptor-positive breast adenocarcinoma MCF-7 cell line (#HTB-22, ATCC) was cultivated in Dulbecco’s Minimum Essential Medium (DMEM, #D6429, Sigma-Aldrich), supplemented with 10% fetal bovine serum (FBS, #F7524, Sigma-Aldrich) and 0.01 mg/mL human recombinant insulin. Triple-negative breast cancer cell line MDA-MB-231 (#HTB-26, ATCC) was cultivated in Dulbecco’s Modified Eagle Medium (DMEM, #D6429, Sigma-Aldrich) with a 10% addition of FBS. Murine triple-negative breast adenocarcinoma cell line 4T1 (#CRL-2539, ATCC) was expanded in Roswell Park Memorial Institute 1640 (RPMI 1640, #61870-010, Gibco) medium with 10% FBS added. All culture media were supplemented with 100 U/mL penicillin/streptomycin (#15140-122, ThermoFisher) and kept at 37° C in a 95% humidified and 5% CO^2^ atmosphere.

### Aptamers and buffers

6-Carboxyfluorescein (FAM) or biotin labelled or unlabelled ssDNA random aptamer library (RND) containing constant primer binding regions and 40 nt randomised region (5’-FAM/Cy5-ATCCAGAGTGACGCAGCANNNNNNNNNNNNNNNNNNNNNNNNNNNNNNNNNNNNNNNNTGGACACGGTGGCTTAGT-3’) and FAM, Cy5 or biotin labelled or unlabelled ssDNA aptamer GreenB1 containing primer binding constant regions and 40 nt sequence in-between (5’-FAM/Cy5-ATCCAGAGTGACGCAGCATGGGGGTAGTGGTGGTTAGGAGTGGAGGCGAGGAGAGCGGTGGACACGGTGGCTTAGT-3′) were purchased from Integrated DNA Technologies (IDT). Oligonucleotides were diluted to 100 µM concentration using DNase and RNase-free water. Aptamers were folded at 10 µM or 1 µM concentration in folding or binding buffer at 95 °C for 5 min, then cooled down to room temperature (RT) for at least 15 min. Binding buffer contained 5 mM MgCl_2_, 4.5 mg/mL D-glucose, 0.1 mg/mL baker’s yeast tRNA and 1 mg/mL bovine serum albumin (BSA) (#A9647, Sigma-Aldrich) in MgCl_2_ and CaCl_2_-free Phosphate Buffered Saline (PBS) (#D8537, Sigma-Aldrich, contained K^+^ at 4.45 mM, Na^+^ at 157 mM concentrations). The folding buffer contained 5 mM MgCl_2_ in PBS. NUPACK was used to predict the secondary structure of GreenB1 (43).

### Aptamer binding to MCF-7 and MDA-MB-231 cells

FAM-RND and FAM-GreenB1 were folded at 1 µM in binding buffer and diluted further (500, 250, 125, 62, 31, 16, 8 nM for MCF-7 and 500, 125, 25, 5 nM for MDA-MB-231). MCF-7 and MDA-MB-231 cells were cultivated in T75 flask (Sarstedt) until 80% confluence. Cells were washed with PBS and dissociated using non-enzymatic cell-dissociation buffer CellStripper (#25-056-CI, Corning) for 5-9 min, followed by the addition of complete culture medium, centrifugation at 300g for 5 min and removal of the supernatant. Cells were washed twice with binding buffer, split into respective samples, and resuspended with different concentrations of FAM-RND or FAM-GreenB1. Samples were incubated on ice for 1 h, washed twice with washing buffer, resuspended in 40 µL of binding buffer and analysed using Amnis ImageStream^X^ Mk II imaging flow cytometer and IDEAS software (Luminex).

### Surface β1-integrin availability after GreenB1 binding *in vitro*

FAM-RND or FAM-GreenB1 aptamers were folded and incubated at 0, 50, 100 and 500 nM concentrations with MDA-MB-231 cells grown on a 6-well plate at 37 °C in an incubator for 24 h. Cells were removed from a 6-well plate using a non-enzymatic cell dissociation solution and incubated on ice with Cy5-GreenB1 aptamer at 100 nM concentration for 1 h. Cells were washed with binding buffer twice, resuspended in 30 µL of binding buffer and analysed using Amnis ImageStream^X^ Mk II imaging flow cytometer and IDEAS software (Luminex).

### Pulse-chase incubation and lysosome co-localisation

MDA-MB-231 cells were cultivated in a 6-well plate until reaching 80% confluence. The Cy5-GreenB1 aptamer was folded in folding buffer at 10 µM concentration and diluted in 1 mL complete growth media to 100 nM before adding to cells. Cells were incubated with Cy5-GreenB1 for 1 h and replaced with complete growth media; afterwards removed for further processing using a non-enzymatic cell dissociation reagent. Cells were analysed 1, 2, 3, 4, and 24 h after adding Cy5-GreenB1. After dissociation, cells were washed twice with PBS/0.1%BSA, resuspended in 100 µL PBS/0.1%BSA and kept on ice. Before imaging flow cytometry, cells were centrifuged at 300 g for 5 min and resuspended in 30 µL of 75 nM LysoTracker green (#L7526, ThermoFisher). Samples were analysed using Amnis ImageStream^X^ Mk II imaging flow cytometer (Luminex).

### Proximity labelling of GreenB1 target protein

Tyramide-AlexaFluor555 working solution was prepared by combining 50 µL of 20X reaction buffer (Component C3 from #B40933),1000 µL purified water, 10 µL of Tyramide-AlexaFluor555 reagent (Component C1 from #B40933) and 10 µL of 0.15% hydrogen peroxide. The tyramide-biotin working solution was prepared by combining 50 µL of 20X reaction buffer (Component C3 from #B40933), 1000 µL purified water, 10 µL of 0.15% hydrogen peroxide and adding Tyramide-Biotin (#LS-3500, Iris Biotech) at 500 µM final concentration.

GreenB1-biotin, unlabelled GreenB1, RND-biotin, and unlabelled RND oligonucleotides were diluted in 500 µL folding buffer to 1 µM concentration and folded as described in the previous section. Horseradish (HRP)-conjugated streptavidin (Component B, 500 µL) from Alexa Fluor 555 Tyramide SuperBoost Kit (#B40933, ThermoFisher) was added to folded oligonucleotides and incubated at RT for 30 min to create oligonucleotide-biotin-streptavidin-HRP complex. The mixture was transferred to a 100 kDa MWCO Amicon Ultra-4 centrifugal filter unit (#UFC810008, Merck) centrifuged at 7500g, refilled four times to remove the unbound aptamer and finally concentrated to approximately 100 µL. The resulting complex was diluted to 500 µL and added to cells for performing confocal microscopy or labelled protein pull-down using magnetic streptavidin beads (#65001, ThermoFisher) afterwards. For confocal microscopy, MDA-MB-231 cells were cultivated in an 8-well culture slide (#354118, Falcon). Cells were washed twice with PBS before applying oligonucleotide-biotin-streptavidin-HRP complexes, followed by incubation at 37 °C for 1 h. The medium was aspirated, and cells were washed 3 times with folding buffer before adding 100 µL of Tyramide-AlexaFluor555 working solution to each well. The reaction was stopped after 2 min by adding 100 µL of 1X stop reagent (100 µL of Component D in DMSO from #B40933 and 1100 µL of PBS) to each well. Cells were washed 3 times with PBS, fixed with 4% formaldehyde at RT for 10 min, and washed twice with PBS. Nuclei were stained with DAPI (#D1306, ThermoFisher) at RT for 5 min and washed once with PBS. Chambers were removed from the slide, and mounting media and coverslip were added to the slide and imaged. For the pull-down experiment, MDA-MB-231 cells were cultivated in a T75 flask, washed with PBS once and dissociated using non-enzymatic cell-dissociation buffer CellStripper (Corning) for 5-9 min, followed by the addition of a complete culture medium. The cell suspension was split into a necessary number of samples (approximately 1*10^6^ cells per sample), centrifugated for 5 min at 300g and supernatant was removed. Cells were resuspended in 500 µL of folding buffer and 500 µL of oligonucleotide-biotin-streptavidin-HRP complexes, followed by incubation at RT for 1 h in an end-over-end rotator. Cells were then centrifuged at 300g, washed twice with PBS and resuspended in 100 µL of tyramide-biotin working solution. The reaction was stopped after 2 min by adding 100 µL of 1X stop reagent (100 µL of Component D in DMSO from #B40933 and 1100 µL of PBS) to each sample. Samples were washed with PBS and centrifuged at 300g for 5 min. The samples were either subjected to flow cytometry to confirm biotinylation or lysed for pull-down of biotinylated proteins. For flow cytometry, samples were incubated with Streptavidin-DyLight-488 (#21832, ThermoFisher) at 20 µg/mL in folding buffer for 10 min, washed 3 times with PBS, fixed with 4% formaldehyde, washed 3 times with PBS, resuspended in 200 µL PBS/0.1%BSA and analysed using flow cytometry (BD Accuri C6 Plus). For protein pull-down, samples were lysed by adding 200 uL of sample lysis buffer (50 µL of 4X sample buffer, 20 µL of 10% of n-dodecyl-β-D-maltoside (DDM) from NativePage Sample Prep kit, #BN2008, ThermoFisher, and 130 µL of PBS) and pipetting to solubilise the proteins. Lysed samples were centrifuged at >20 000g at 4 °C for 30 min. The supernatant was collected and added to 20 µL of Dynabeads MyOne Streptavidin C1 beads (#65001, ThermoFisher) per sample. The lysate was incubated with magnetic beads at RT on an end-to-end rotator for 30 min. Beads were washed four times with 200 µL of sample lysis buffer. Beads were transferred to a new 1.5 mL centrifuge tube after each wash. Elution was achieved by adding 30 µL of 25 mM biotin in lysis buffer and heating at 95 °C for 5 min. Biotin elution strategy was adapted from Cheah and Yamada (44). Elution was repeated two times, and the supernatant was collected. The third elution step was done by adding 30 µL of reducing sample buffer and heating at 95 °C for 5 min.

### Tris/Glycine gel electrophoresis and mass spectrometry

Reducing sample-loading buffer (2 µL) was added to 10 µL of each elution from streptavidin beads after proximity labelling. Samples were heated at 95 °C for 5 min and loaded on 12% Mini-PROTEAN TGX precast protein gel (#4561043, Bio-Rad). The gel was run using 1x Tris/Glycine running buffer at 100 V for 90 min. The gel was stained using the SilverQuest Silver staining kit (#LC6070, ThermoFisher). The bands of interest were cut out and sent for mass spectrometry proteomics analysis at the University of Tartu Proteomics core facilities. (https://www.tuit.ut.ee/en/research/proteomics-core-facility). Figure 3d was prepared using R (45) in RStudio(version 2021.9.2.382) (46) and packages *readxl* (47), *ggplot2* (48) and *ggrepel* (49).

### Electrophoretic mobility shift assay with α3β1-integrin

GreenB1 or RND aptamers were folded at 1 µM concentration and diluted to 35 nM using a folding buffer with 5% glycerol. α3β1-integrin protein complex (#2840-A3-050, R&D Systems, 20 µL) at 360 nM (100 µg/mL) in PBS was diluted to 180 nM (molar concentrations calculated based on sodium dodecyl-sulfate polyacrylamide gel electrophoresis SDS-PAGE migration of each protein under reducing conditions, 150 kDa for α3-integrin and 125 kDa for β1-integrin) using a folding buffer with 10% glycerol. Further dilutions were prepared using a folding buffer with 5% glycerol. To each 10 µL of α3β1-integrin dilutions, 10 µL of 35 nM of GreenB1 or RND were added. Final concentration for aptamers was 17.5 nM and α3β1-integrin concentrations were 90, 45, 22.5, 11.25, 5.61, 2.8, 1.4 and 0.7 nM. The mixture was then incubated at RT for 2 h and loaded on 3% agarose gel prepared using 0.5x Tris/Boric acid buffer without ethylenediaminetetraacetic acid (EDTA) and run at 180 V in a cold room (4 °C) using 0.5x Tris/Boric acid as running buffer for 30 min. The gel was stained with SYBR Gold nucleic acid stain (#S11494, ThermoFisher) in 0.5x Tris/Boric acid buffer for 30 min and destained in purified water for 10 min. K_d_ was calculated from triplicate measurements using Prism 9.3.1 (GraphPad) using one site-specific binding equation (Y = Bmax*X/(Kd + X)).

### Fluorescence polarization (FP) assay with α3β1-integrin

FP reactions (25 μL) were set up in black, non-transparent flat-bottom 384-well microplates (#3821, Corning). Each reaction contained PBS supplemented with 5 mM MgCl_2_, 0.05% Tween-20, 10 nM of FAM-labelled GreenB1 aptamer or FAM-labelled RND, and varying concentrations of β1-integrin (#10587-H08H1, SinoBiological). Reactions were performed in triplicate. Measurements were recorded on a Hidex Sense microplate reader equipped with 485nm/535nm optical filters, using 100 flashes, medium lamp power and a PMT voltage of 750 V. Titration data were fitted in Prism 9.3.1 (GraphPad) using the following equation:

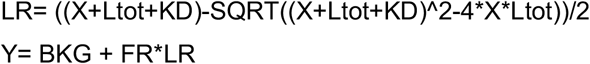

Where:

X - concentration of integrin-β1 (serial 2-fold dilutions)

Ltot - total concentration of aptamer (fixed)

PR - fluorescence ratio, a unitless constant (fitted)

BKG - background polarisation of unbo und aptamer (fitted)

KD - dissociation constant (fitted)

Y - fluorescence polarisation (recorded)

### Aptamer homing *in vivo* and tissue imaging

Prior to aptamer homing *in vivo* analysis in the 4T1 cell TNBC orthotopic model, binding of FAM-GreenB1 and FAM-RND to 4T1 cells at 100 nM was confirmed using DB Accuri C6 Plus flow cytometer. Animal experimentation protocols were approved by a Committee of Animal Experimentation of the Estonian Ministry of Agriculture (Permit #159). FAM-GreenB1 or FAM-RND (2 nmol) were injected i.v. to orthotopic 4T1 tumour-bearing mice (n=3 in each group). The mice were anaesthetised, perfused with 20 ml PBS, and tumour, heart, liver, lung, and kidney were collected after 1 h of circulation. Organs were flash-frozen in liquid nitrogen and kept at -80 °C until further use. Organs were sectioned at 30 µm and fixed in methanol (equilibrated at -20 °C) for 2 min. 5% BSA in PBS with 0.05% Tween-20 was applied to tissue sections for 30 min at RT to block non-specific antibody binding. FAM-GreenB1 or FAM-RND was detected using a 1:200 dilution of rabbit anti-fluorescein antibody (#A-11090, ThermoFisher) at RT for 1 h, followed by 1:400 dilution of secondary anti-rabbit AlexaFluor555 labelled antibody (#A32794, ThermoFisher) at RT for 1 h. Staining with secondary antibody alone was used as a negative control. Five random fields of view were imaged from all organs from all mice. Both primary and secondary antibody labelled, and secondary antibody alone were imaged by fluorescent microscopy. For statistical analysis, mean pixel intensity from primary and secondary antibody labelled organ sections were divided by mean pixel intensity from the corresponding organ from the same animal when stained with secondary antibody alone. Nuclei were stained using Hoechst 33342 (#62249, ThermoFisher) solution at 20 µM concentration. Fluorescence microscopy was done using Nikon C2 microscope (Japan) and images were analysed using Nis-Elements C 4.13 software. Fluorescence confocal microscopy was done using Olympus FV1200MPE, Germany. Whole tumour imaging was performed using slide scanner Aperio VERSA Brightfield, Fluorescence and FISH Digital Pathology Scanner, USA. Statistical analysis was done in Prism 9.3.1 (GraphPad) using 2-way ANOVA followed by Šidák’s multiple comparison test. Figure 6b was prepared using R (45) in RStudio (46) and packages *readxl* (47), *ggplot2* (48), *ggsignif* (50).

## RESULTS

### GreenB1 binds to cultured triple-negative breast cancer cells

GreenB1 aptamer was originally identified by us in a SELEX on cultured malignant cells (51). Whereas GreenB1 was identified in a screen for clear cell renal cell carcinoma cell line binders, it also selectively bound MDA-MB-231 breast cancer cell line. FAM-labelled GreenB1 aptamer (Fig. 1A) or FAM-labelled 40 nt randomized oligonucleotide library (RND) were incubated with human TNBC cell line MDA-MB-231 and progesterone and estrogen receptor-positive human breast cancer cell line MCF-7. Incubation was performed in the presence of increasing concentrations (5–1000 nM) of either GreenB1 or RND on ice for 1 h. Cell-bound fluorescence was analysed using imaging flow cytometry. GreenB1 resulted in significantly increased fluorescence intensity (Fig. 1C and Fig. 1F) at 5 nM concentration after the incubation with the TNBC cell line MDA-MB-231, whereas only a moderate fluorescence increase was observed when incubated with the MCF-7 cell line (Fig. 1E and Fig. 1G). In contrast, RND aptamer library did not show increased binding over the range of concentrations to MDA-MB-231 (Fig. 1B) and MCF-7 (Fig. 1D) cell lines. These results suggest that the GreenB1 binds selectively to cultured TNBC cells.

**Figure 1.**
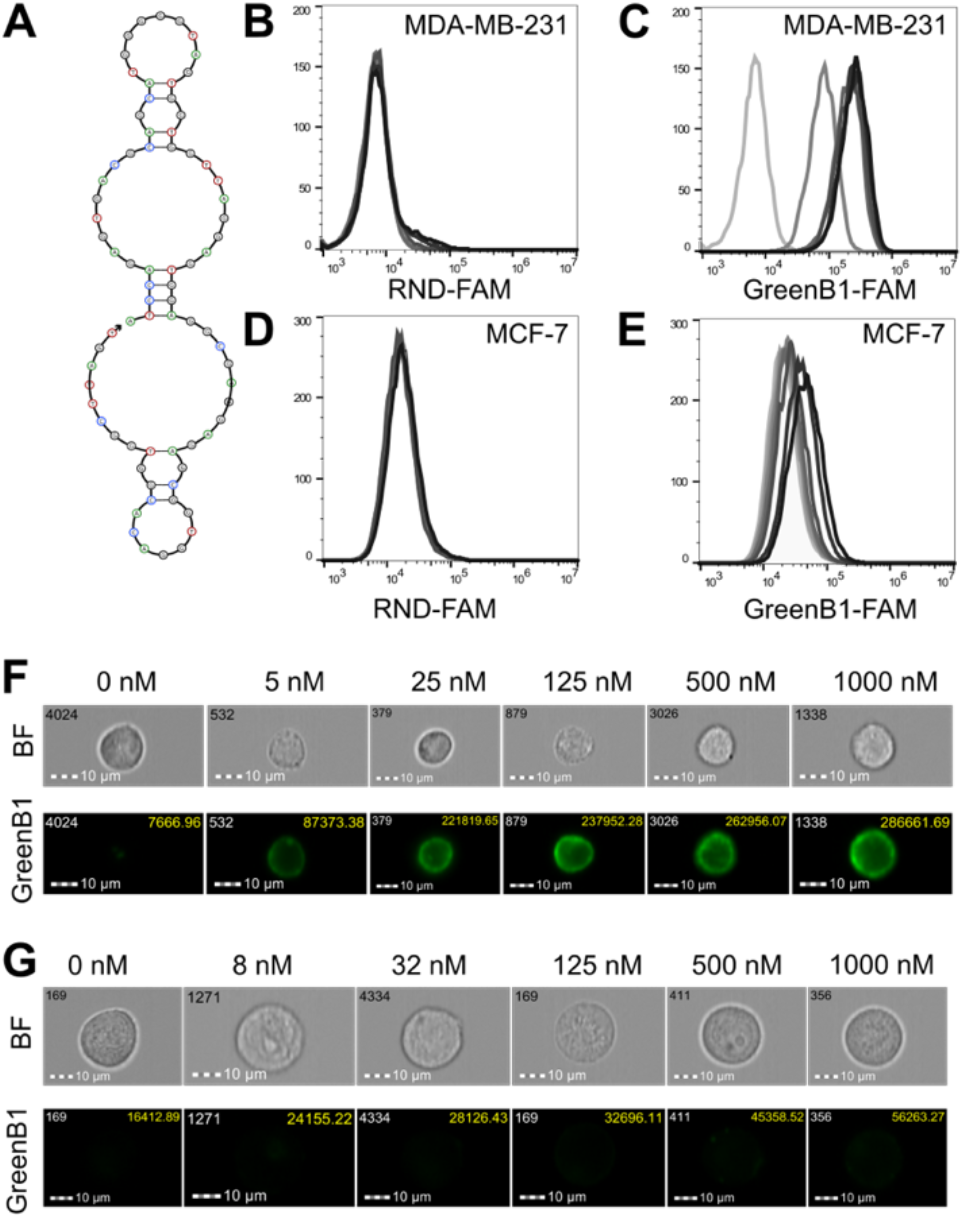
GreenB1 binds to triple-negative breast cancer cells. GreenB1 structure as predicted by NUPACK (A). Random aptamer (B) and GreenB1 (C) binding to MDA-MB-231 triple-negative breast cancer cell line at 0, 5, 25, 125, 250, 500, 1000 nM concentrations. Random aptamer (D) and GreenB1 (E) binding to MCF-7 progesterone and estrogen receptor-expressing cell line at 0, 8, 16, 32, 62, 125, 500, 1000 nM concentrations. Imaging flow cytometry images of GreenB1 binding to MDA-MB-231 (F) and MCF-7 (G) cell lines at different concentrations. The values in the right upper corner of the FAM channel image show the fluorescence intensities of the whole cells.

### Proximity labelling identifies β1-integrin as the target protein

GreenB1 target protein was identified by a proximity ligation-based approach (Fig. 2). GreenB1-biotin or RND-biotin were complexed with streptavidin-horseradish peroxidase (HRP) and incubated with live MDA-MB-231 cells, followed by a proximity labelling reaction using tyramide-AlexaFluor555 or tyramide-biotin.

**Figure 2.**
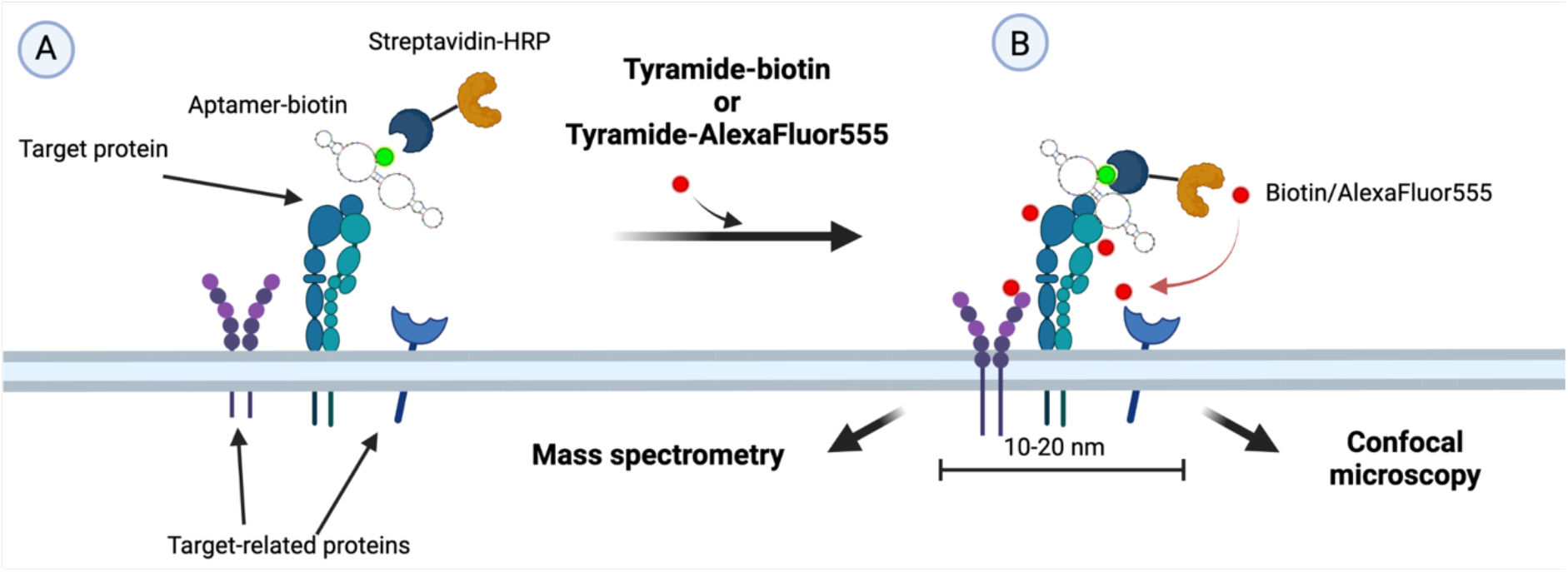
GreenB1 protein target identification using proximity labelling. Biotin labelled GreenB1 or RND was incubated with horseradish peroxidase (HRP) conjugated to streptavidin. Complexes or streptavidin-HRP alone were incubated with live MDA-MB-231 cells for 1 h (A). After washing away the unbound complex, tyramide-biotin or tyramide-AlexaFluor555 with hydrogen peroxide was added to cells for 2 min. HRP, in the presence of hydrogen peroxide, creates a highly reactive tyramide species that labels nearby proteins (B). Fluorescently labelled proteins were further imaged using confocal microscopy. Biotinylated proteins were pulled down using streptavidin-coated magnetic beads, eluted using 25 mM biotin in lysis buffer and heating at 95 °C for 5 min. Eluates were run on the gel and further analysed using mass spectrometry.

The reaction of GreenB1-HRP complex with tyramide-AlexaFluor555 on MDA-MB-231 cells resulted in staining observable under a confocal microscope (Fig. 3A) with a much higher intensity than when using the RND complex (Fig. 3B).

**Figure 3.**
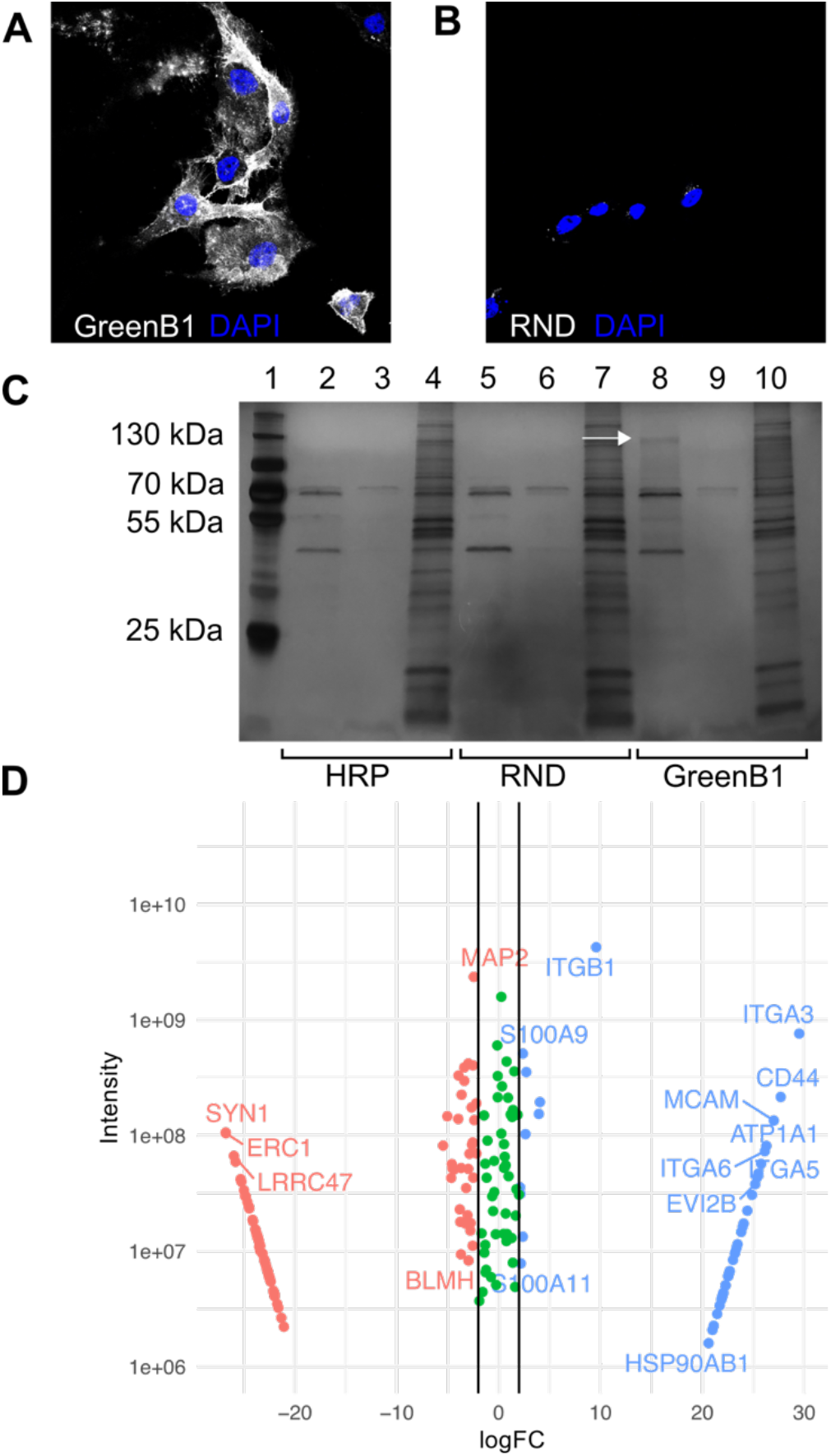
Proximity labelling results for GreenB1 target identification. Confocal microscopy images of proximity labelling using Tyramide-AlexaFluor555 with MDA-MB-231 cells after binding of GreenB1-HRP complex (A) or RND-HRP complex (B). Pull down results from proximity labelling with tyramide-biotin. Lane 1 contains a marker. Streptavidin-HRP (lanes 2, 3, 4), RND-HRP (lanes 5, 6, 7) and GreenB1-HRP (lanes 8, 9, 10). The first two lanes in each sample were eluted using biotin and heat; the third was eluted using reducing sample buffer (C). Mass spectrometry proteomics of control and target bands corresponding to 130 kDa. Log2 fold change on the x-axis. Combined signal intensity from both samples on the y axis (D).

Biotinylated proteins were pulled down using streptavidin-covered magnetic beads and eluted using excess biotin and heating, followed by elution using reducing sample buffer and heating. Eluted proteins were run on Tris/Glycine gel and subjected to silver staining. A single band of ∼130 kDa was observed in the first eluate of the GreenB1 sample (Fig. 3C, lane 8) but not in RND (Fig. 3C, lane 5) or streptavidin-HRP alone (Fig. 3C, lane 2) samples. The region containing the band (Fig. 3C, indicated with a white arrow) in the GreenB1 sample and the corresponding molecular weight region from the RND sample were subjected to mass-spectrometry (MS) proteomics analysis. Two proteins with the highest MS intensity and highest logarithmic fold-change (logFC) difference between RND and GreenB1 samples were the β1 and α3 integrins. Molecular weight of β1 integrin (around 120-130 kDa) (52) on SDS-PAGE and the location of target band further supported the β1 integrin being the target protein for the GreenB1 aptamer.

### GreenB1 has a K_d_ value against β1-integrin in the low nanomolar range

Mass spectrometry resulted in several additional hits besides α3 and β1 integrins (Fig. 3D). To confirm the binding and determine the dissociation constant (K_d_) of GreenB1 for α3β1 and β1 integrins, we used electrophoretic mobility shift assay (EMSA) and fluorescence polarization (FP) analysis. For EMSA, GreenB1 or RND at 17 nM concentration were incubated with α3β1-integrin at increasing concentrations and separated by electrophoresis on 3% agarose gel. Whereas GreenB1 band decreased in intensity with increasing α3β1-integrin concentration (Fig. 4A), no change was seen for RND (Fig. 4B). The calculated K_d_ for GreenB/α3β1-integrin interaction was 15 nM (95% CI 8 – 26 nM) (Fig. 4C).

**Figure 4.**
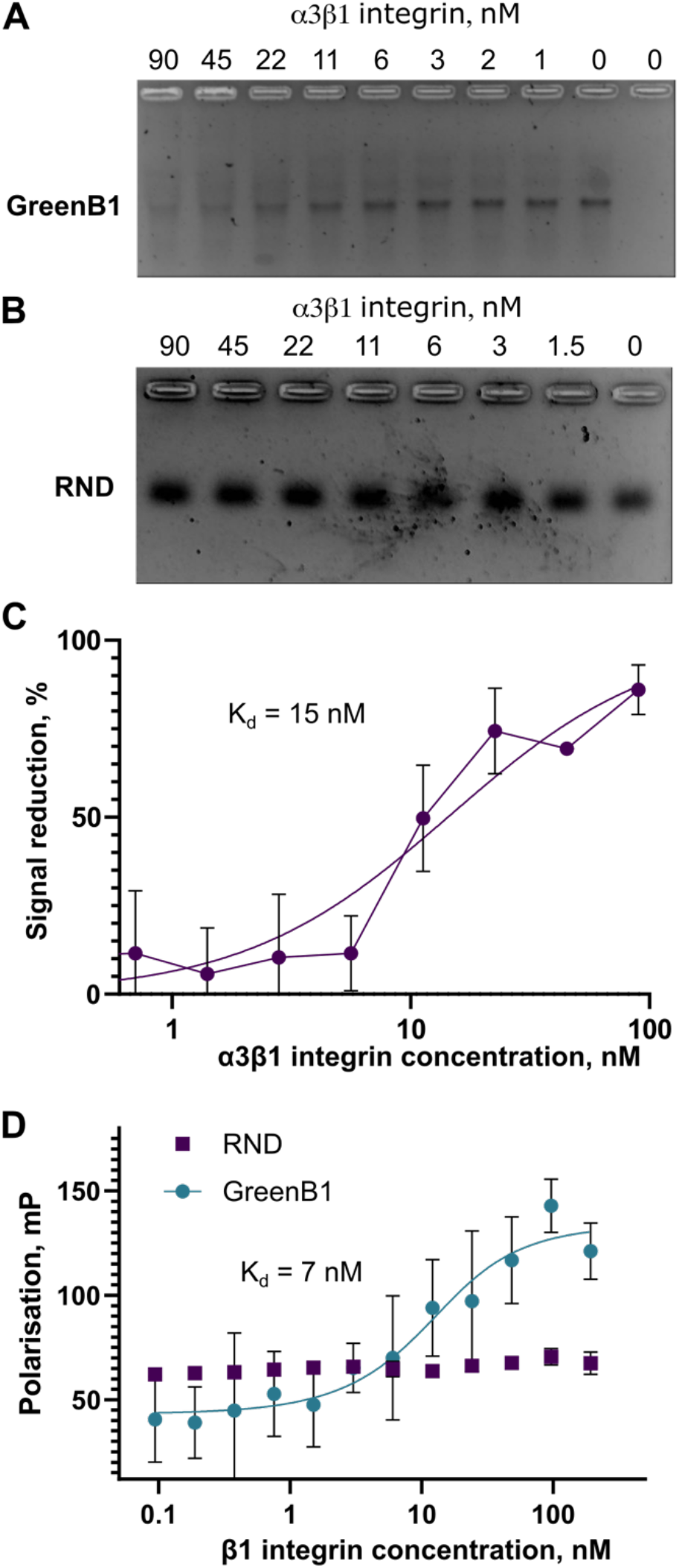
GreenB1 binding to α3β1-complex and β1-integrin. Electrophoretic mobility shift assay (EMSA) using increasing concentrations of α3β1-integrin protein at 17.5 nM fixed concentration of GreenB1 (A) or RND (B) aptamers. GreenB1 and α3β1-integrin EMSA results plotted as reduction of GreenB1 signal intensity (K_d_ = 15 nM, 95% CI 8-26 nM) (C). Fluorescence polarization using 10 nM of FAM-labelled GreenB1 aptamer or FAM-labelled RND and varying concentrations of β1-integrin. (K_d_ = 7 nM, 95% CI 0 – 17 nM) (D). Plots depict averages of triplicate measurements ± SE and the fitted model.

GreenB1 binding to β1-integrin alone was further tested using fluorescence polarization (FP). β1-integrin had a much higher signal intensity than α3-integrin in MS proteomics results, suggesting that it could be the target protein within α3β1-integrin complex. Varying concentrations of β1-integrin were incubated with 10 nM of FAM-labelled GreenB1 or FAM-labelled RND. An increase in FP was observed for GreenB1 but not for RND (Fig. 4D). K_d_ = 7 nM (95% CI 0 – 17 nM) was calculated using Prism 9.3.1 (GraphPad). The results from EMSA and FP show that GreenB1 aptamer binds to β1 integrin in the low nM range.

### GreenB1 target protein is available for binding after the pre-incubation with GreenB1

To find out whether GreenB1 binding to the β1-integrin has an impact on the β1-integrin density on the cell surface, we tested GreenB1 binding dynamics by pre-incubating variable concentrations of either FAM-RND or FAM-GreenB1 with MDA-MB-231 cells for 24 h. After the pre-incubation, FAM-RND/FAM-GreenB1 containing medium was replaced with fresh medium and both samples were incubated with 100 nM Cy5-GreenB1. FAM-RND pre-incubation did not show concentration-dependent FAM fluorescence increase (Fig. 5A), and Cy5-GreenB1 was binding to the target protein afterwards (Fig. 5B). FAM-GreenB1 pre-incubation resulted in a concentration-dependent increase in FAM fluorescence (Fig. 5C). However, Cy5-GreenB1 binding was not affected (Fig 5D). Lack of changes indicated that target protein remained available for aptamer binding irrespectively of the FAM-GreenB1 concentration during pre-incubation.

**Figure 5.**
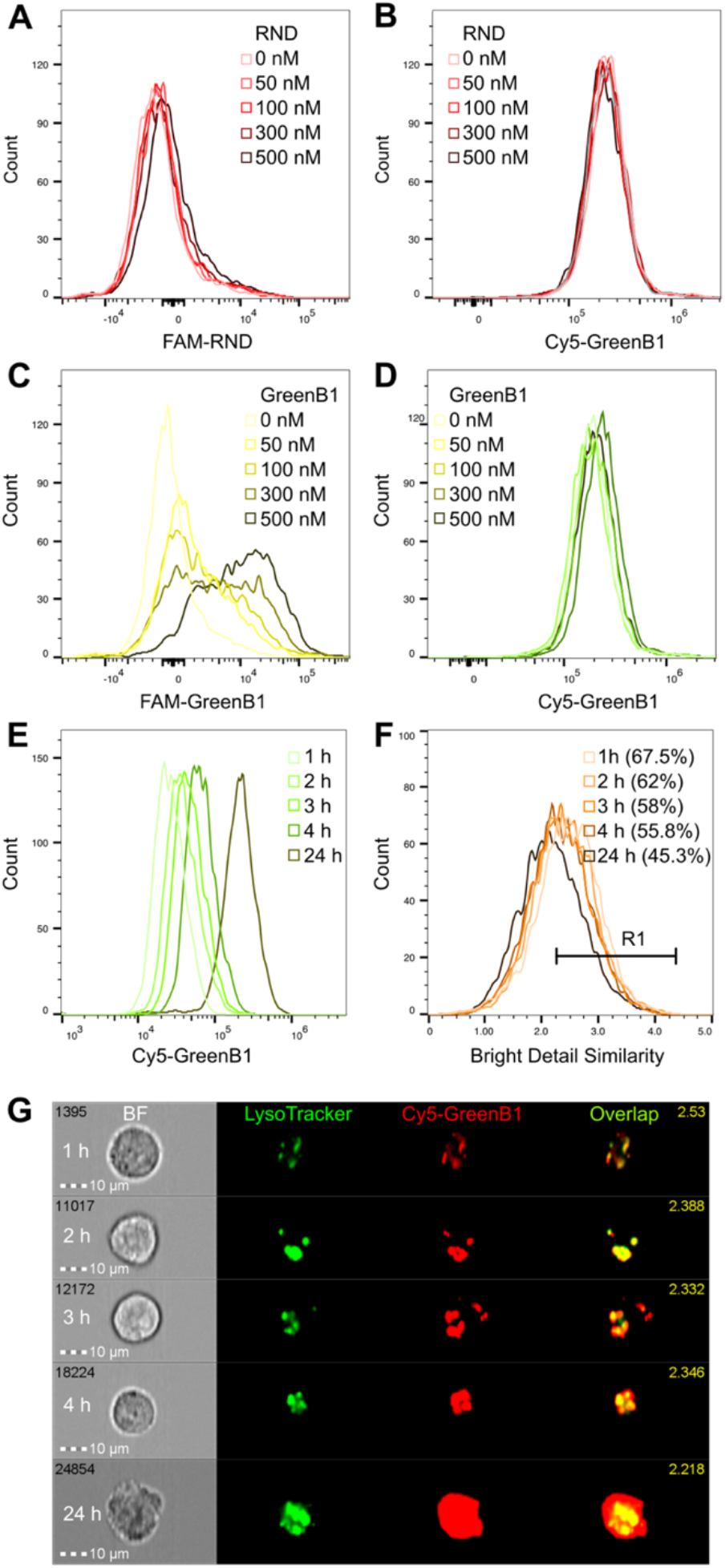
Cellular internalization cycle of GreenB1 aptamer. FAM-labelled RND aptamer library pre-incubation with MDA-MB-231 cells on a 6-well plate at different concentrations for 24 h (A) followed by incubation with 100 nM Cy5-GreenB1 at 4°C for 1 h (B) shows no concentration-dependant fluorescence increase. FAM-labelled GreenB1 aptamer pre-incubation for 24 h with MDA-MB-231 cells on a 6-well plate at different concentrations (C), followed by 100 nM Cy5-GreenB1 incubation at 4°C for 1 h (D) results in concentration-dependant fluorescence increase during the pre-incubation but does not affect Cy5-GreenB1 binding afterwards. Cy5-labelled GreenB1 aptamer pulse-chase incubation 100 nM with MDA-MB-231 cells in a 6-well plate results in time-dependent Cy5-fluorescence intensity increase (E). Cy5-GreenB1 co-localization with LysoTracker Green shows the highest co-localization based on bright detail similarity between LysoTracker Green and Cy5-GreenB1 at 1 h after incubation and a slight decrease after 24 h (F). Representative images of Cy5-GreenB1 co-localization with lysosomes after different time points (G)

**Figure 6.**
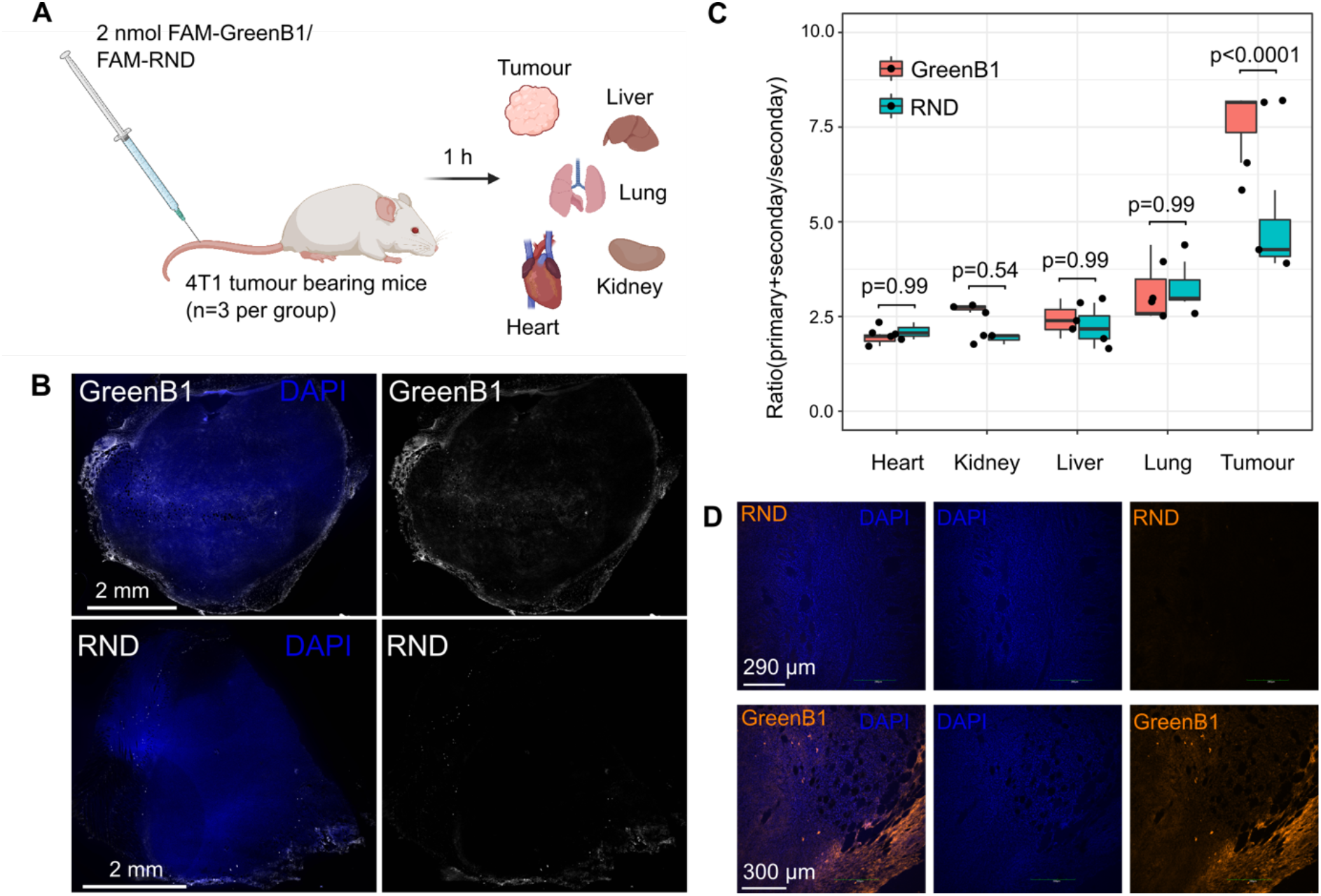
*In vivo* homing of systemically-administered GreenB1 aptamer. FAM-GreenB1 or FAM-RND were injected i.v. into Balb/C mice (n=3 per group) carrying orthotopic murine 4T1 triple-negative breast cancer. The tumour and control organs were collected (A). Whole tumour imaging from mouse carrying 4T1 tumour when injected with FAM-GreenB1 or FAM-RND and labelled with primary anti-FAM antibody and secondary antibody (B, first row for GreenB1, second row for RND). Mean fluorescence intensity from each organ was calculated from five fields of view. A statistically significant fluorescence intensity difference was detected between tumours from mice injected with FAM-GreenB1 compared to those from mice injected with FAM-RND (p<0.0001). There were no statistically significant fluorescence differences between control organs (heart, kidney, liver, lung) from FAM-GreenB1 injected mice and FAM-RND injected mice (C). Representative confocal microscopy images of tumours labelled with anti-FAM antibody and secondary antibody with AlexaFluor555 from mice injected with FAM-GreenB1 (D, second row) or FAM-RND (D, first row).

### GreenB1 rapidly internalises in cells and shows co-localisation with lysosomes

The aptamer uptake was further studied using pulse-chase fluorescence imaging. Cy5-labelled GreenB1 was incubated with MDA-MB-231 cells in a complete culture medium at 100 nM for 1 h. The cells were collected immediately, or aptamer-containing medium was replaced with a fresh culture medium without the aptamer and incubated for additional 2, 3, 4 and 24 h. 75 nM LysoTracker Green DND-26 was added to each sample before imaging flow cytometry. Co-localization analysis was done using IDEAS software based on bright detail similarity in both fluorescence channels. On the one hand, Cy5-GreenB1 signal intensity increased with each time point, indicating fast initial binding to the membrane-bound target protein and slower subsequent internalization (Fig. 5E). However, bright detail similarity was highest after 1 h (67.5% of cells were determined to be co-localized events) and it decreased with each following time point (2 h = 62%, 3 h = 58%, 4 h = 55.8%, 24 h = 45.3%), indicating trafficking to endolysosomal pathway (Fig. 5F and Fig. 5G).

### Systemically administered GreenB1 homes to TNBC lesions modelled in mice

We next tested the ability of GreenB1 to home to murine 4T1 TNBC (53) *in vivo*. We first confirmed that GreenB1 binds to cultured 4T1 cells (Supplementary Figure S1). Mice bearing orthotopic 4T1 triple-negative breast cancer lesions were injected with either 2 nmol of FAM-GreenB1 or FAM-RND i.v. (Fig. 6A). Imaging of whole tumour cryosections confirmed higher accumulation of FAM-GreenB1 within the tumour (Fig. 6B, first row for tumour stained with both anti-FAM primary antibody and secondary antibody, secondary antibody alone as Supplementary Figure S2A) compared to FAM-RND (Fig. 6B, second row for tumour stained with both anti-FAM primary antibody and secondary antibody, secondary antibody alone as shown on Supplementary Figure S2B). GreenB1 had high intensity at the margins of the tumour, but it also reached the tumour core.

Statistically significant fluorescence difference was detected in 4T1 tumour (p<0.0001), indicating preferential homing of GreenB1 to triple-negative breast cancer model in mice (Fig. 6C). In contrast, GreenB1 showed no statistically significant fluorescence differences in control organs when compared against RND (p=0.99 for heart, p=0.54 for kidney, p=0.99 for liver and p=0.99 for lung). Representative confocal images showed higher FAM-GreenB1 signal intensity within the tumour (Fig. 6D, second row) compared to the FAM-RND signal (Fig. 6D, first row).

## DISCUSSION

TNBC is the most lethal of the breast cancer subtypes with the estimated median overall survival time for metastatic TNBC from 10 to 13 months. However, when detected early, at stage I, TNBC has a 5-year survival rate of 85%, which is lower than other breast cancer subtypes (54). Chemotherapy in combination with ICIs has been demonstrated to improve median progression-free survival and median overall survival in PD-L1 positive subgroups (12, 55), but an ideal therapeutic approach remains elusive (56). Precision-guided interventions hold promise for delivering therapeutic agents to tumours and aptamers with high selectivity towards target molecules are promising candidates for targeted therapy or diagnostics purposes. We demonstrate that at low nanomolar concentrations, aptamer GreenB1 selectively binds to the triple-negative breast cancer cell line MDA-MB-231 but not to the estrogen- and progesterone receptor-expressing breast cancer cell line MCF-7. On cell surface, GreenB1 interacts with β1-integrins. GreenB1 is subsequently internalized via the endolysosomal uptake pathway, while β1-integrin is recycled back to the cell surface. *In vivo*, systemically administered GreenB1 aptamer homes to 4T1 cell triple-negative breast cancer lesions modelled in mice.

It is likely that GreenB1-based targeting strategies can be used for precision delivery of drugs and imaging agents to the β1-integrin-positive solid tumours other than TNBC. Integrins have been extensively used in cancer therapeutic affinity targeting efforts. Several antibody-based integrin αV and αVβ3/β1/β5 targeting therapies have been tested in Phase I/II clinical trials with disappointing results (56). Antibody delivery to poorly vascularized tumour tissue could be hampered by their large molecular weight. Smaller-sized iRGD peptide targeting αVβ3/β5 integrins has shown promising preclinical results for pancreatic ductal adenocarcinoma therapy (57) and is now investigated in Phase II clinical trial (ClinicalTrials.gov Identifier: NCT03517176). The NRP-1 binding CendR motif in iRGD promotes extravasation into the tumour (58). GreenB1 is suited for development into an affinity targeting ligand capable of accessing difficult-to-reach tumour vasculature since aptamers are several orders of magnitude smaller than antibodies.

GreenB1 co-localization with lysosomes suggests that it is internalized in cells and likely being trafficked via the route previously established for oligonucleotide delivery. According to it, GreenB1 dissociates from β1-integrin in early endosomes due to a pH shift that causes an aptamer conformation change. GreenB1 is then transported to lysosomes whereas β1-integrin follows the re-cycling pathway (59).

Lysosome-targeting chimaeras (LYTACs) have taken advantage of lysosome shuttling proteins to target membrane-bound and extracellular proteins for degradation and could be used to act on currently “undruggable” proteins (60). Bispecific aptamer-based LYTAC system used insulin-like growth factor type II receptor (IGF-IIR) as lysosome shuttling component to degrade targeted proteins (61). GreenB1 trafficking to lysosomes implies that more research on the application of this aptamer to produce LYTACs that work via the β1-integrin re-cycling route is warranted. Furthermore, it is likely that by modifying GreenB1-based targeting to allow endolysosomal escape, the system can be adapted for delivery of payloads into the cytosol and other intracellular compartments. This will be particularly important for large siRNA and peptide cargoes with polar and charged character that are unable to translocate efficiently into the cytosol to exhibit their biological activity (59, 62).

Unmodified aptamers have a circulation half-life of minutes to hours and are degraded in serum by exonucleases. The circulating half-life of GreenB1 can be extended by adding high molecular weight compounds such as poly-ethylene glycol (PEG), creating multivalent constructs larger than the glomerular filtration rate cut off (50-60 kDa) (15, 63) or circularising the aptamer to make it less susceptible to nuclease digestion (64). GreenB1 can be further linked to cytotoxic chemicals via a lysosome-sensitive linker (65) or liposomes containing an anticancer payload (66) to assess its ability to diminish tumour burden.

In addition to applications in targeted delivery, GreenB1 may have inherent functional activity due to modulating the status of its target integrins. Integrins are known to profoundly regulate cell migration, survival, and proliferation. Compared to many cell-surface proteins that are degraded or do not change their location after ligand binding, integrins are constantly trafficked and recycled within cells (67). Integrin expression modulation is linked to cancer invasion, formation of metastatic lesions, tumour growth and development of resistance to treatment (68). In breast cancer, receptor tyrosine kinase c-Met can replace α5 integrin as β1-integrin binding partner, forming a complex that drives cancer cell migration due to higher affinity to fibronectin (69). In TNBC, β1-integrin and Talin-1 (TLN1) interaction blocking has recently been described as a potential therapeutic target (70). Silencing of β1-integrin has been shown to increase the sensitivity to cancer drugs and inhibit cancer cell migration and invasion (71). β1-integrin is a required protein for forming vasculogenic mimicry, a tumour blood-supply mechanism where cancer cells form blood vessel-like structures (72). GreenB1 has a high affinity for β1-integrin, suggesting that it could be used therapeutically to disrupt β1-integrin interactions with TLN1 or c-Met, hence altering TNBC cell invasiveness. Alternatively, research into a GreenB1-based strategy that silences β1-integrin activities in TNBC and thus increases susceptibility to existing therapies is necessary.

As a technical advancement, we adopted a proximity labelling-based approach (widely used to study protein-RNA/DNA and protein-protein interactions) to identify GreenB1 engagement partners. Compared to extract-based techniques, such as affinity chromatography, the proximity ligation has the advantage of yielding less background and more relevant hits. To generate reactive species from a substrate, biotin ligases or peroxidases connected to a targeting moiety, such as an aptamer, are used. The activated substrate then covalently bonds to neighbouring proteins and can be utilized to pull down proteins close to the binding point (73). Our unpublished studies show that a similar approach can be used to identify binding partners for other targeting ligands, such as peptides, and that the technique can be even applied for *in vivo* interaction studies (Haugas et al., manuscript in preparation).

In summary, here we report a new β1-integrin binding aptamer GreenB1 that selectively binds TNBC cells *in vitro*, is quickly internalized in cells, and *in vivo* preferentially homes to the TNBC lesions in mice. GreenB1 translational applications are of great interest in the future and might lead to innovative targeted protein breakdown or therapeutic approaches.

## DATA AVAILABILITY

Fluorescent microscopy images used for statistical analysis have been submitted to BioImage Archive (74) and are available here: https://www.ebi.ac.uk/biostudies/studies/S-BIAD483.

Fluorescent microscopy images used for statistical analysis (.nd2 format under “GreenB1-in-vivo-images”), confocal images (.oib format under “GreenB1-confocal-images”), slide scanner images (.scn and .sdf formats under “GreenB1-slide-scanner-images”), imaging flow cytometry files (.rif, .cif, .daf and .cpm formats under “GreenB1-RND-MCF7-MDA-MB-231-different-conc”) and mass spectrometry proteomics (.raw and .xlsx data table under “greenb1-ms-proteomics”) have been deposited on BioStudies (75) and available here: https://www.ebi.ac.uk/biostudies/studies/S-BSST857.

The mass spectrometry proteomics data have been deposited to the ProteomeXchange Consortium (76) via the PRIDE (77) partner repository with the dataset identifier PXD034982 and 10.6019/PXD034982.

## SUPPLEMENTARY DATA

Supplementary Figures are available at NAR Online.

## ACKNOWLEDGEMENT

We would like to thank Toomas Jagomäe for whole-tumour imaging with slide scanner and University of Tartu Proteomics core facilities for mass spectrometry proteomics analysis. Figure 2 and Figure 6A were created using BioRender.com.

## Author contributions

K.P. developed and carried out the proximity ligation, imaging flow cytometry, EMSA, immunostaining and confocal microscopy experiments, as well as the statistical analyses.M.H. developed initial proximity ligation protocol, carried out in *vivo* experiments and collected confocal microscopy images. T.P. and E.P. designed, carried out, and analysed the fluorescence polarisation experiment. V.P. performed the cryosectioning and immunostaining. T.T., U.R., and K.P. conceived and oversaw the project, and wrote the manuscript. The final manuscript was read and approved by all authors.

