## Supplementary Figures for "Targeting triple-negative breast cancer with β1-integrin binding aptamer"

### β1-integrin binding and lysosome targeting aptamer homes to triple-negative breast cancer in mice

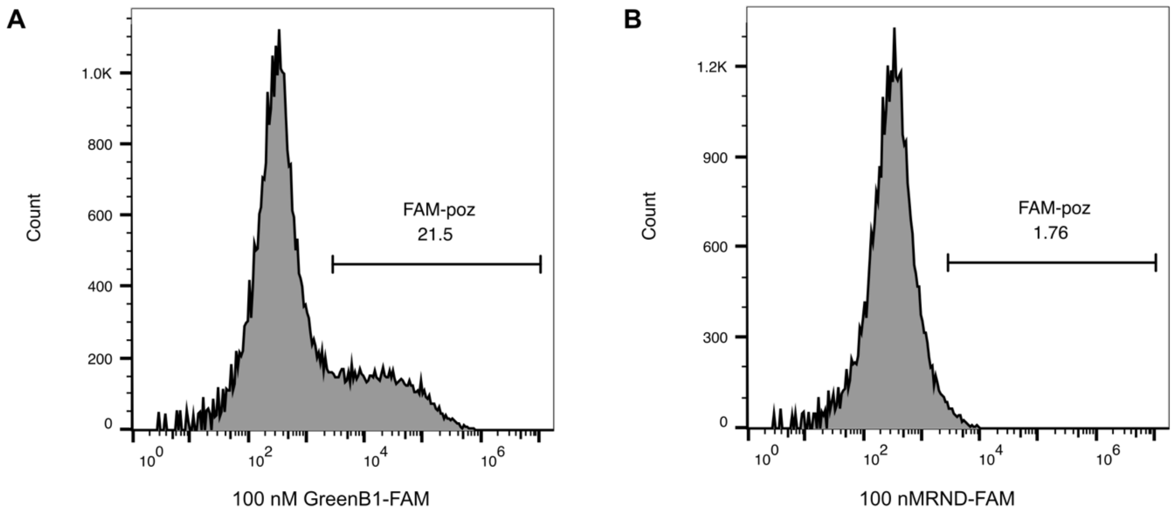


**Supplementary Figure S1. GreenB1 aptamer binding to 4T1 cells.** FAM-GreenB1 or FAM-RND were folded and diluted to 100 nM concentration using binding buffer. 4T1 cells were detached using non-enzymatic cell dissociation buffer, washed, and incubated with FAM-GreenB1 or FAM-RND for 1 hour on ice. Cells were then washed two times and analysed using BD Accuri C6 Plus. Higher fraction of 4T1 FAM⁺ cells in GreenB1 sample (A) compared to RND sample (B) confirm GreenB1 binding to 4T1 cells.

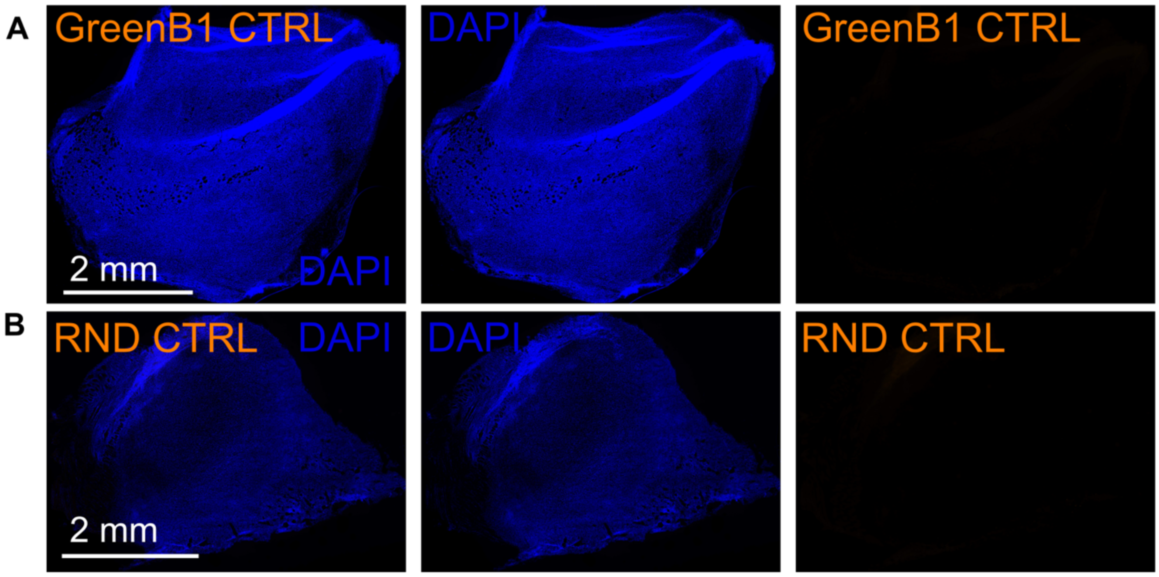

**Supplementary Figure S2. GreenB1 homing to tumour lesions *in vivo*.** Whole tumour imaging from mouse carrying 4T1 tumour when injected with FAM-GreenB1 (A) or FAM-RND (B) and labelled with only secondary antibody.
